# The First 1000 Days: An Agent-Based Model of Early Language Acquisition

**DOI:** 10.64898/2026.03.28.715023

**Authors:** Hadas Raviv, Arkadii Tsyhanov, Kira Gousios, Aja Altenhof, Haocheng Wang, Berlin Chen, Ofri Raviv, Tal Rosenwein, Casey Lew-Williams, Liat Hasenfratz, Uri Hasson

## Abstract

A longstanding challenge in developmental science is to understand how children learn language from naturalistic everyday input. To study this process, we leveraged the First 1,000 Days (1kD) dataset, which provides longitudinal, ultra-dense daily audiovisual recordings for individual children in their home environments. This unusually detailed, child-specific record of early experience enabled us to pair each child’s rich language input with a cognitively grounded learning agent, linking naturalistic experience (“nurture”) to internal learning mechanisms (“nature”). Trained incrementally on each child’s input without prior linguistic knowledge, the learning agent discovered speech units corresponding to the English phoneme inventory and acquired thousands of words, closely mirroring individual developmental trajectories. Learning generalized across children while preserving individual differences in rate and timing. Interestingly, learning relied not only on linguistic input but also on the rehearsal of past experiences at the end of each training day. These findings demonstrate that everyday environments provide sufficient structure for language acquisition and establish a unified mechanistic framework for studying development in real-world contexts.

## Introduction

Understanding how children acquire language has long been a central question in psychology, linguistics, and philosophy^1–6^. Children learn language in complex, dynamic, and highly individualized environments, yet they reliably converge on a shared functional use of linguistic units. They acquire different components of language on distinct timescales (e.g., phonemes, words, grammar), largely without explicit instruction. Nativist theories argue that the speed, efficiency, and robustness of acquisition depend on innate structures or representations that encode key linguistic knowledge^2,3,7,8^. In contrast, emergentist and usage-based accounts contend that children’s environments provide rich, structured linguistic and contextual input, from which speech units and grammatical patterns can arise through domain-general learning mechanisms^4–6,9^. Despite decades of research, a fully explicit mechanistic account remains lacking - one that links an individual infant’s day-to-day experience from birth to the neural processes that shape development over time, while capturing the coupled dynamics across multiple levels of language (phonemes, words, and grammar).

The First 1,000 Days (1kD) project addresses this challenge directly by integrating ultra-dense, daily recordings of individual children’s linguistic input (nurture) with a cognitively grounded learning agent (nature) within a **unified framework**. We ask: how do internal learning mechanisms interact with naturalistic language input as language development unfolds over time in individual children? As a starting point, we focus on two of the earliest learning problems - phoneme learning and word learning - which typically unfold within the first two years of life^10–16^. To that end, we (1) measure children’s everyday language environments using ultra-dense longitudinal recordings, and (2) pair these recordings with a cognitively plausible learning agent that acquires phonemes and words by processing the speech stream experienced by an individual child. Crucially, our goal is to simulate learning processes on realistic developmental timescales, using each child’s ecologically rich input. We evaluate our learning agent by comparing its learning dynamics to the developmental trajectories observed in individual children.

### Capturing Nurture

Dense, ecologically valid recordings are crucial for developing learning agents that acquire language under input conditions that approximate children’s real-world experiences. Over the last few decades, the field has increasingly moved toward naturalistic, child-centered data collection. Recent advances - including daylong audio, ambient sensing, egocentric video, and wearable devices ^17–21^ - have greatly expanded researchers’ ability to document children’s everyday environments. Datasets such as SAYCam^22^, BabyView^21^, and corpora of day-long audio recordings^23^ have also increased the ecological and demographic diversity of available recordings. However, most existing datasets still provide only brief snapshots of a child’s development, covering only a few hours or days. As a result, they do not capture the temporal details of children’s learning trajectories, nor do they provide the density required to train models that learn exclusively from individual children’s input patterns.

Capturing the full scope of early linguistic environments is ethically, technically, and linguistically challenging; it requires a fundamental rethinking of every step of the research process. The 1kD project addressed this challenge, inspired by Deb Roy’s pioneering Speechome project^24^, but enabled by modern advances in data collection and machine learning^25^. We undertook a large-scale replication and expansion of this approach, leveraging contemporary audiovisual sensors, cloud infrastructure, and automated processing tools. We recorded 12–14 hours of naturalistic audiovisual data per day from 15 racially and economically diverse U.S. families across the first 1,000 days of each child’s life^25^. From this collected corpus, we analysed and modeled a subset of eight infants, which will be the focus of this paper.

All data were securely stored in a dedicated research environment in the AWS cloud and processed via an automated pipeline that analyzed millions of one-minute files to detect the child’s location and extract all surrounding speech. Across 495–990 days of recording days per child (median 960), we obtained 3–7.5 hours of high-quality speech input **per day**. This yielded an unprecedented quantitative characterization of the *environmental input* available to each child, providing the necessary data density to observe how idiosyncratic experience shapes language acquisition.

### Learning From Nurture

With these dense recordings in hand, we built a learning agent to simulate how individual children process speech and learn words from their own everyday linguistic input. The model is intentionally minimalist: it relies solely on self-supervised learning objectives, is initialized randomly, and has no prior knowledge of phonemes, word boundaries, or household-specific vocabulary. Instead, learning is driven solely by each child’s recorded input and proceeds in a developmentally plausible, incremental way, with day-by-day updating.

Deep learning frameworks, including modern large language models (LLMs), provide a powerful statistical approach to discovering structure in high-dimensional data and have achieved striking performance on many language tasks^26–28^. However, current industrial training practices, relying on hundreds of billions of text tokens and repeated exposure to the same datasets, bear little resemblance to the conditions of human development^29^. A child encounters only a tiny fraction of the input used to train contemporary LLMs (Gilkerson et al., 2017; Bergelson et al., 2023).^30,31^ Moreover, children’s experience is embedded in spoken, multimodal, socially grounded interaction, rather than in large-scale, decontextualized text corpora^5,6,32^. To overcome these limitations, recent research has focused on training models with smaller or more naturalistic datasets^33–41^. However, two key limitations remain. First, prior models rely primarily on sparsely sampled child-centered corpora, which do not capture the longitudinal structure and variability of children’s everyday linguistic experiences. Second, training protocols often depend on cognitively unrealistic assumptions - such as innate tokenizers or discrete textual units - which do not reflect the continuous, noisy acoustic stream children actually encounter^37^.

Our learning agent is based on deep learning architectures and is designed to model development under ecologically plausible conditions, addressing key gaps in prior work. Specifically, it incorporates: (1) ultra-dense, child-centered input from the 1kD project; (2) individualized training for each child’s unique linguistic environment; (3) a shared agent architecture that must account for learning across all children; (4) continuous, day-by-day training with daily replay, inspired by memory consolidation processes^42^; (5) a curriculum aligned with broad stages of language development^43^; and (6) a fully learned- rather than pre-specified lexicon.

Using this framework, our cognitively feasible learning agent learned discrete speech units directly from continuous household audio. Over a developmentally equivalent period of one year, these emergent units became increasingly aligned with phonetic structure while remaining fine-grained and context-sensitive. This representation supported the gradual emergence of phoneme-level regularities and the acquisition of thousands of words from each child’s input stream. The agent’s lexicon growth closely tracked the trajectory of the corresponding child, as measured by standardized behavioral assessments (MacArthur–Bates Communicative Development Inventories; CDI^44^). A key ingredient in reproducing these developmental trajectories was a daily replay: each day, the agent reprocesses a limited amount of previously heard speech (“replay amount”), consistent with the idea that temporal structure and memory processes shape learning^42^. Critically, the same learning architecture generalized across all eight households.

Individual differences in learning rate were captured by three internal parameters: replay amount, updates per hour, and learning rate, which together are interpretable as learner-internal variation (nature) interacting with differences in experience (nurture).

The combination of dense 1kD recordings and the learning-agent framework offers, to our knowledge, the first mechanistic account of how incremental learning processes interact with the structure of infants’ everyday language environments to support early language acquisition as it unfolds in real-life contexts.

## Results

We combined ultra-dense longitudinal recordings of children’s first 1,000 days with a novel learning-agent approach to examine how children acquire speech sounds and build vocabulary in their home-language environments. Below, we describe each step in detail and present the resulting empirical analyses, along with comparisons between the model and the children.

### Section 1: Ultra-dense 1,000-day recording of language developments at home

#### Data Collection and Processing

With human subjects approval (Princeton University IRB, USA; protocol #13884), a data safety monitoring board, and parental consent, we installed audiovisual recording systems in fifteen homes to capture the daily experiences of infants across the first 1,000 days of life (Fig. 1-A). The participating households were racially and economically diverse, reflecting U.S. Census demographics. Recordings were conducted 12-14 hours per day in key household spaces using one to two cameras with built-in microphones, along with one high-fidelity external microphone per room. To process the approximately 1.2 million hours of data, we developed a scalable, AI-based processing infrastructure on AWS and a multimodal analysis pipeline designed to detect speech occurring near each infant. Full details of the recording setup and processing pipeline are provided in Raviv et al.^25^.

**Figure 1.**
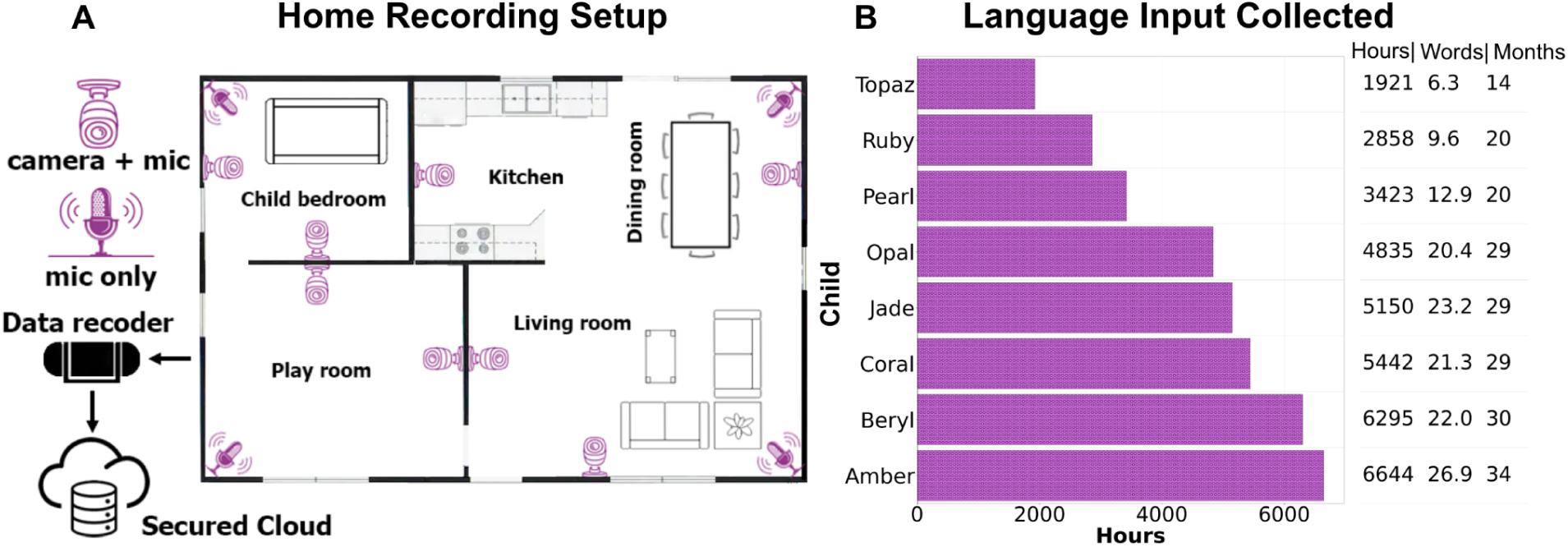
Overview of in-home recording setup and data collection. (A) Example layout of recording equipment across key rooms in a participating home. (B) The linguistic input collected across the eight families: total hours with child and speech presence, total number of words (in Millions), and total months for training.

We have completed the recordings, preprocessing, and modeling phases for data from eight homes, which will serve as the primary focus of this paper. The recording in eight homes generated approximately 820 hours of audiovisual data per day. Families were recorded for a median of 960 days (range: 495–990 days). The analysis process yielded approximately 1,900–6,600 hours of naturalistic audio per child (Fig. 1-B, all children’s names are pseudonyms), which were automatically transcribed, producing corpora totaling roughly 6-27 million words. We further segmented the recordings into daily “awake-time” speech hours using each child’s daily schedule (Section 2; Methods) - resulting in timelines spanning 14 to 34 months across infants.

The naturalistic far-field recordings include background environmental noise and reverberation. Our goal is to model how children learn from the speech present in their environment. Therefore, we simplified the input for this study by resynthesizing a clean, speech-only signal from the transcripts using a text-to-speech (TTS) model^45^. Full details of the processing pipeline are provided in the Methods section.

#### Quantifying infants’ word acquisition

In addition to the continuous audiovisual recordings, we collected monthly caregiver-reported measures of children’s language development using the MacArthur–Bates Communicative Development Inventories (CDI^44^). From 8 to 16 months, we administered the Words & Gestures form to assess comprehension and production. At 16 months, following standard practice, we transitioned to the Words & Sentences (Toddler) form to track speech production through the end of the study. Full details of the CDI collection and analysis are provided in the Methods and in Raviv et al.^25^.

Word-learning trajectories varied noticeably among children, with the age at which half of CDI words were acquired ranging from 12 to 22 months (Fig. 2). To derive these trajectories, we defined the age of acquisition for each lexical item as the first month in which the caregiver reported that the child knew the word. The cumulative number of learned words as a function of age (in months) is shown as green (comprehension) and blue (production) points in Fig. 2-A. We fit a sigmoid function to each trajectory, fixing the asymptote (maximum CDI vocabulary size) and estimating two parameters: Time to Learn (**TL**), the age at which 50% of the words had been acquired, and steepness (**S**), which reflects the rate of learning. Enabled by dense longitudinal sampling, this model provided the best fit to individual trajectories^46,47^. The fitted parameters for comprehension trajectories across the eight infants are shown in Figs. 2-B and 2-C. These analyses reveal inter-individual differences in word-learning trajectories that are visible only at the level of densely sampled individual data. Notably, variability in Time to Learn (Fig. 2-B) exceeded variability in steepness (Fig. 2-C), suggesting larger differences in *when* children reached mid-vocabulary than in *how quickly* vocabulary grew once acquisition began.

**Figure 2.**
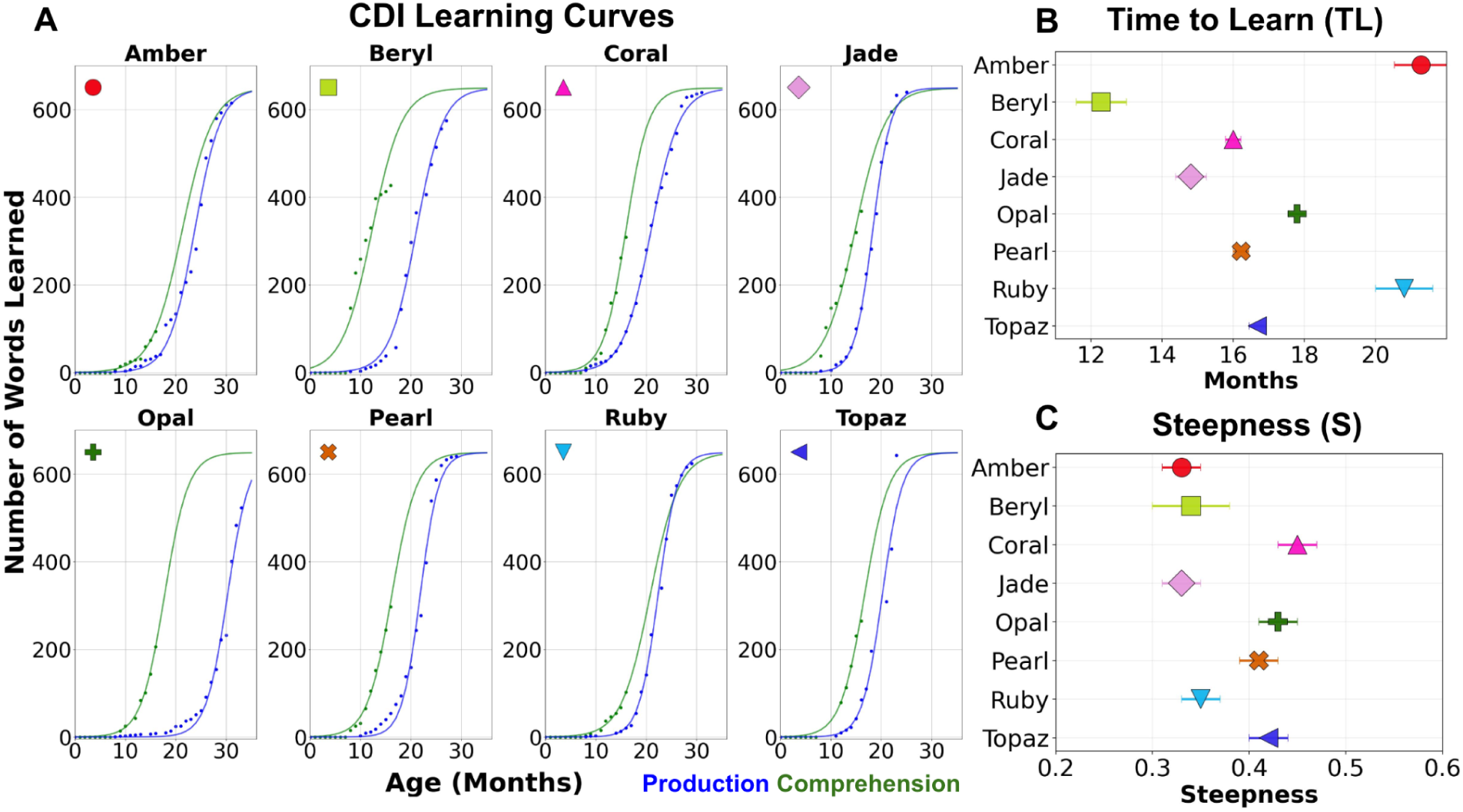
Word learning trajectories derived from parental reports. (A) Data points show the cumulative number of words learned as a function of age (in months) for comprehension (green) and production (blue), based on parental responses to the CDI. Solid lines indicate fitted sigmoid curves for word comprehension and production for each of the eight infants. (B) Fitted values of **TL**, the age at which 50% of the words were acquired, for the comprehension curves shown in panel A. (C) Fitted values of **S**, the steepness of the curve reflecting learning rate, for the comprehension curves shown in panel A.

### Section 2: Designing a Cognitively Feasible Learning Agent

The ultra-dense recordings enabled us to build a learning agent that can model how everyday experiences interact with internal processes over time, gradually supporting the acquisition of phonemes and words. Importantly, we designed a self-supervised learning agent that relies solely on speech input available to each child, without using any prior knowledge of speech or language structure. The training protocol is designed to be cognitively feasible, supporting gradual day-by-day learning, curricular learning, and memory replay during sleep. In the following sections, we describe the model architecture and learning protocol in detail.

#### Learning Agent Architecture

Our learning agent prioritizes cognitive plausibility by employing an architecture that learns directly from speech input in a fully unsupervised manner and continuously updates its representations through exposure. The architecture follows an encoder–decoder design, decomposing learning into two stages. First, the encoder learns to discretize the continuous speech stream into 20-ms speech units^38,40,43,48^ analogous to phonemes or subphonemic elements. Second, once the encoder has stabilized, the decoder updates its weights to learn sequential regularities by predicting the next upcoming discrete unit given preceding context. As the decoder’s predictive performance improves, sequences of units (e.g., subwords, words, short phrases) that become reliably predictable are treated as learned tokens.

To operationalize this cumulative process, we introduce a third component, the “dictionary”, which incrementally stores learned sequences as learning progresses (Fig. 3-A). For the encoder, we used DINO-SR^49^, which learns cluster-centered representations of auditory space via a data-driven, unsupervised objective. For the decoder, we used a speech-tailored Transformer, partially adapted from Kharitonov et al.^50^. Full architectural details and hyperparameters are provided in the Methods section.

**Figure 3.**
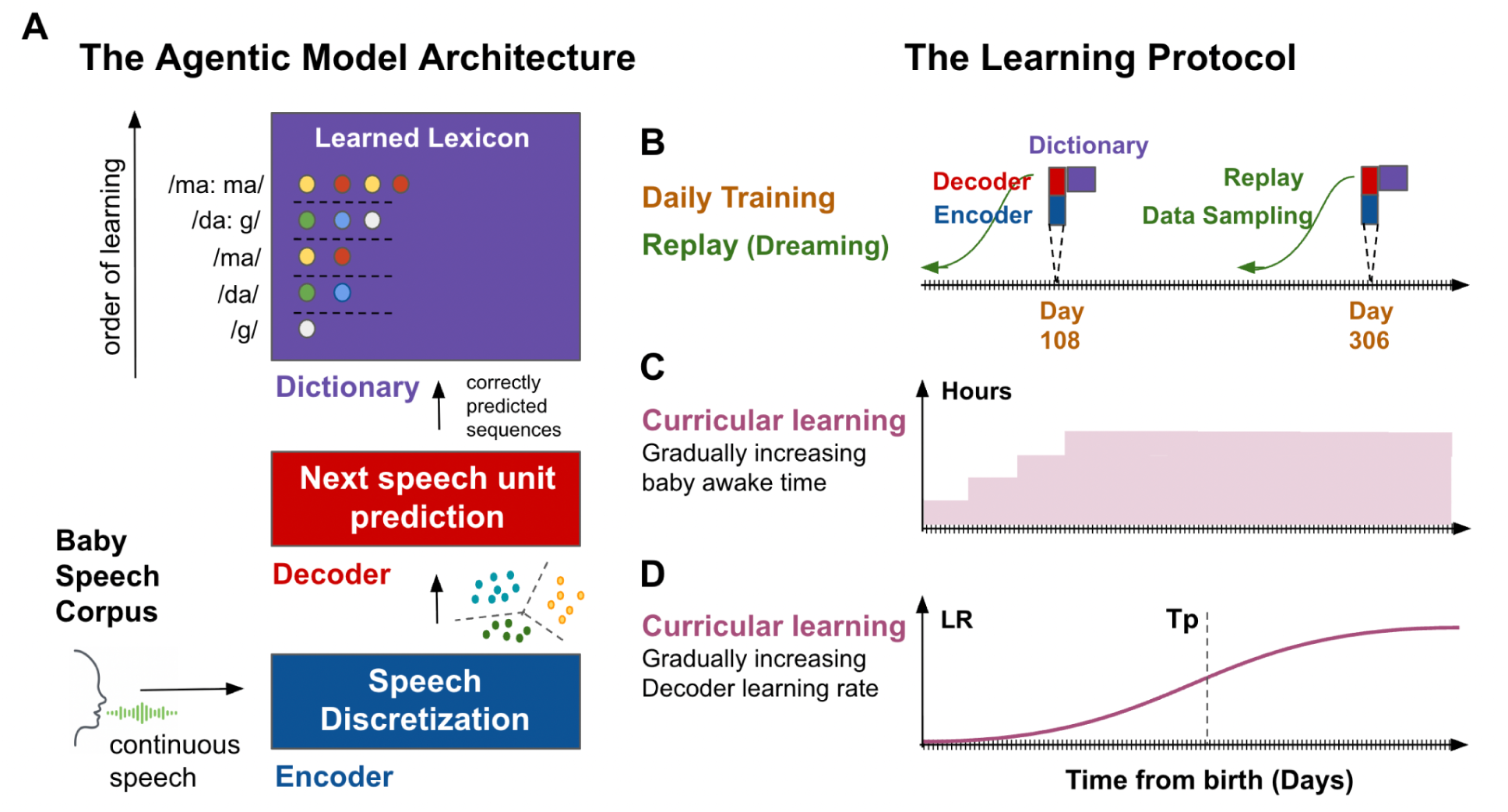
Overview of the model architecture and training protocol. (A) The agentic model comprises three modules. The encoder is trained on continuous speech from the infant speech corpus and learns a mapping from continuous speech to discrete speech units. The decoder receives distinct units and their durations and learns to predict the next unit. The lexicon aggregates reliably predicted sequences over time. (B-D) Cognitively motivated training protocol: (B) learning from daily amounts of speech estimated directly from the recorded home data; replay, implemented as daily resampling of past experience using a decaying probability with temporal distance; and (C-D) curriculum learning, introduced by gradually increasing infants’ waking hours and by modulating decoder learning via a controlled learning-rate schedule.

#### Training Protocol

We developed a cognitively grounded training protocol characterized by three properties:

1. The agent’s learning trajectory can be explicitly mapped onto real developmental time (days, months, and years). The agent consumes audio in the exact chronological order in which it was recorded, thereby preserving the temporal structure of the input, unlike conventional training regimes that often rely on shuffled data. The basic temporal unit of training is **one day** (Fig. 3-B). For each day, the amount of speech presented to the agent is estimated by multiplying the words per hour derived from household recordings by the number of daily awake hours (see below).
2. Drawing on evidence from human memory replay, we incorporate a biologically inspired **replay** mechanism (Fig. 3-B). In standard deep learning, training is often organized into epochs, with multiple passes over the dataset with examples shuffled across iterations. In our setting, this would be akin to a child repeatedly re-experiencing their first three years of input in a randomized order. Instead, at the end of each day of training, we augment that day’s input with replayed speech segments sampled from earlier days, using a decaying probability function that decreases with temporal distance from the current day (see Methods). This design is motivated by sleep-dependent replay mechanisms implicated in memory consolidation^42^.
3. Learning follows a developmental curriculum that approximates real-world children’s experience. We implement **curriculum learning** in two complementary ways. First, we progressively increase the number of daily awake hours during which the infant is exposed to speech, reflecting real developmental patterns in which early months involve relatively short awake periods (Fig. 3-C). For simplicity, the awake schedule is currently held constant across children. Second, we stage learning across the encoder and decoder to better approximate developmental trajectories (Fig. 3-D). Because acquiring stable speech units (e.g., phonemes or subphonemic elements) is plausibly a prerequisite for learning longer-range temporal regularities (e.g., words), we modulate the decoder’s effective learning rate using a sigmoid schedule. This approach preserves continuous training while attenuating decoder updates early in development, so that learning of sequential patterns becomes more effective only after the encoder has stabilized its speech-unit representations.

Finally, one internal model parameter is batch size, defined as the number of words the model processes before updating its weights. Cognitively, this parameter is analogous to the amount of information accumulated across the processing-timescale hierarchy prior to learning^51^. To account for individual differences in learning pace across children, we fit the learning rate schedule, replay, and batch size separately for each child (Fig. 2B, C). By integrating the measured quantity and content of input with these internal model parameters, the model simulates how experience and learning interact over development on realistic timescales.

#### Evaluating the agent’s learning trajectories

A central challenge in this work is determining which phonemes and words the learning agent has acquired at any given point in development. Given the time-dependent nature of infant learning, in which linguistic elements are encountered intermittently and at varying frequencies, we introduce a weekly evaluation framework to assess the agent’s development. Specifically, we evaluate the agent’s learning at the end of each week using data from the following week, which has not been used for training. This testing protocol mirrors real-world development: both the agent and the child are learning by the ordered sequence of experiences accumulated up to that point, and their knowledge state is assessed on subsequent, previously unseen input.

### Section 3: Training a learning agent based on the linguistic input of a single child

We first present the full training, evaluation, and analysis workflow using data from a single household, focusing on one child (Coral, pseudonym) as an illustrative case study. The learning agent was trained on all child-centered speech from Coral’s home in chronological order, proceeding day by day, with additional resampling (“replay”) of recently encountered input between daily training sessions. Learning was driven by internal self-supervised objectives, and the model was initialized with random weights and had no prior linguistic knowledge. Because the current experiments use clean, resynthesized speech, we allocated the first two months of developmental time (days 0–60) as a placeholder for future work aimed at enabling the agent to separate speech from background sounds in naturalistic audio. In evaluation, we first characterize the development of Coral’s agent speech-unit (sub-phonemes to phonemes) representations and then analyze the emergence of word-level predictions and lexical learning over time. After establishing this end-to-end analysis for Coral’s agent, we train additional agents using ultra-dense recordings from the additional families (seven replications) to evaluate the learning agent’s robustness and generalization across children.

#### Quantifying phoneme acquisition in learned speech units

After 24 months of training, Coral’s agent showed selective associations between learned speech units and 38 of the 39 phonemes (Fig. 4-A). The encoder outputs 512 discrete speech-unit labels at a 20 ms frame rate, providing sub-phonemic temporal resolution. To characterize speech unit-phoneme associations, we computed an association matrix in which each entry represents the probability that speech unit u was assigned to phoneme p during evaluation (Fig. 4-A; see Methods). High (dark) values indicate that a unit is specifically utilized for a particular phoneme. Across 38 phonemes, each phoneme was supported by at least one highly selective unit (with a matrix entry value greater than 0.5), and each speech unit typically exhibited weak associations with other phonemes. For each phoneme, we identified the unit with the highest association value, and the average of these maximum values was 0.91, ranging from 0.53 to 1.00. The only phoneme with a substantially lower value was /ZH/, which was too rare in the data. Overall, these results indicate that the learned units capture fine-grained phonemic structure, with multiple speech units representing each phoneme.

**Figure 4.**
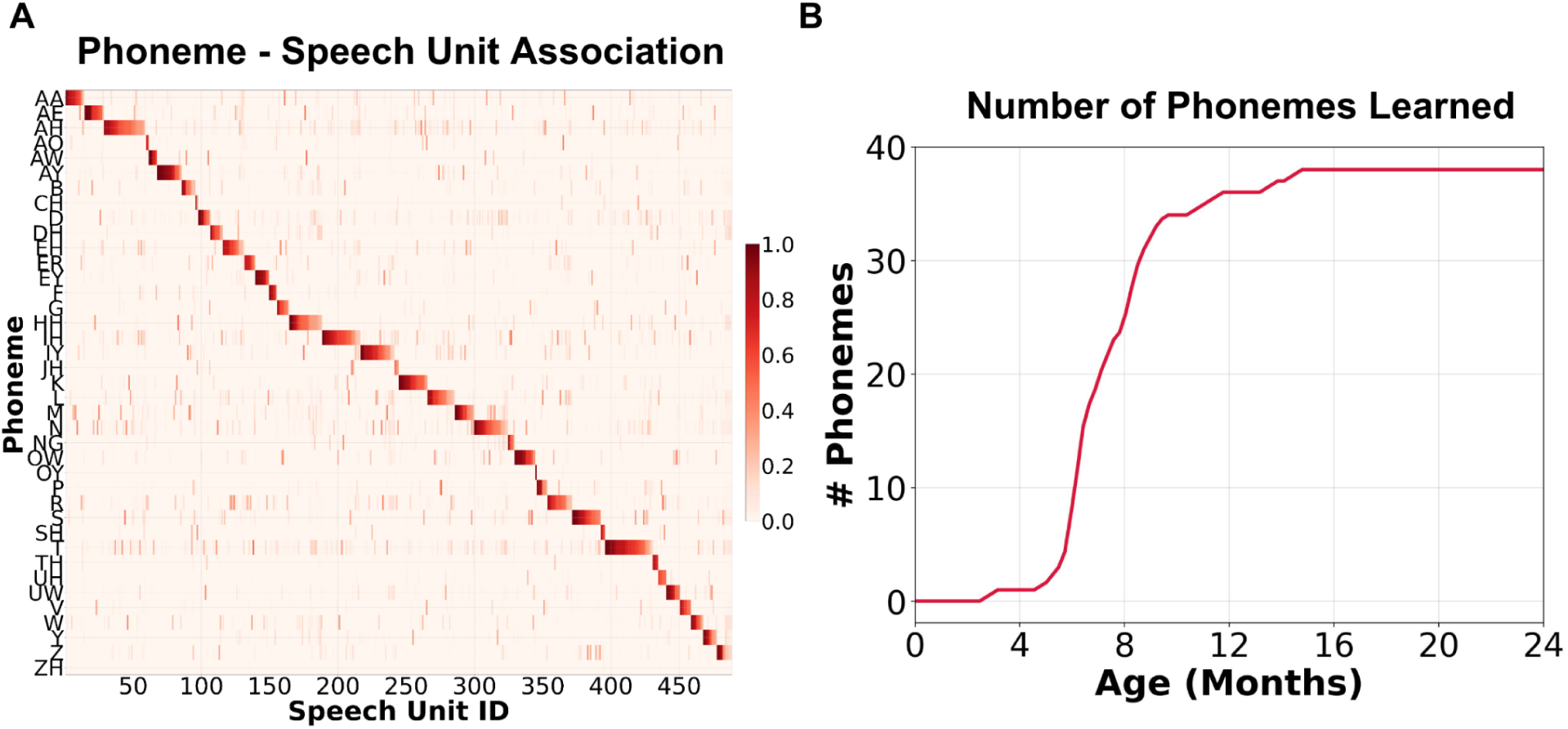
Phoneme - unit structure and phoneme acquisition over time. (A) Phoneme - speech unit association matrix at 24 months for Coral’s agent. (B) Cumulative number of learned phonemes over development using the phoneme-coverage criterion (see Methods).

By 10-12 months of training, Coral’s agent had learned to reliably discriminate most English phonemes. To quantify how this organization emerged over development, we computed the speech unit-phoneme association matrix at the end of each training week and asked when individual phonemes became reliably discriminable. We defined a phoneme as learned when at least 30% of its instances were associated with selective speech units with association values greater than 0.5 (see Methods; Fig. 4-B). As shown, performance rose rapidly over the first five months, reached a high level by the eighth month, and largely stabilized by the sixteenth month. By approximately 15 months, 38 of 39 phonemes had been learned. Additional analyses in the Supplementary materials (Fig. S-1 A-D) further show that speech unit–phoneme mappings developed in a many-to-many structure, preserving fine-grained detail while increasingly consolidating around dominant units associated with each phoneme over development.

#### Quantifying word learning

We next tracked word learning at the decoding stage by evaluating the agent’s ability to accurately predict a sequence of speech units associated with a word in context. In our setting, the decoder predicts each upcoming speech unit as a function of past context. As learning progresses, the decoder confidently predicts sequences of distinct speech units within word boundaries^52,53^. We count a word occurrence as correctly predicted when the agentic model’s confidence remains high (or increases) over a transcript-aligned window spanning at least 90% of the word’s duration (see Methods). To reduce noise, we define a word type as learned at time *t* if, by time *t*, it has been correctly predicted in at least *N* = 15 occurrences across different contexts.

Coral’s agent acquired new vocabulary at a pace that closely matched Coral’s observed development, as reported in monthly CDI assessments (Fig. 5 A-C). We used caregiver reports of word comprehension to estimate the infant’s knowledge of a curated set of 649 CDI words (excluding phrases; see Methods) through 16 months of age (Fig. 5-A, solid green bars). After 16 months, when comprehension data collection ended, we used a sigmoid fit to extrapolate the number of learned words in the same set from 16 to 24 months (Fig. 5-A, striped green bars). At 12 months, Coral understood 97 words (model fit: 91); by 24 months, predicted comprehension reached 632 words (Fig. 5-A). Overall, the agent - like Coral - learned nearly all CDI words at a comparable rate along the developmental trajectory (Fig. 5-B). Over the 24 months, Coral and the agent show similar growth rates (slopes) and learning dynamics for CDI words. Figs. S2–S3 (supplementary materials) report sensitivity analyses over key parameters (N, coverage, mistake score, and skipped units; see Methods), showing that the main results are robust across reasonable settings and that the agent’s trajectories under all conditions closely follow Coral’s developmental trajectory.

**Figure 5.**
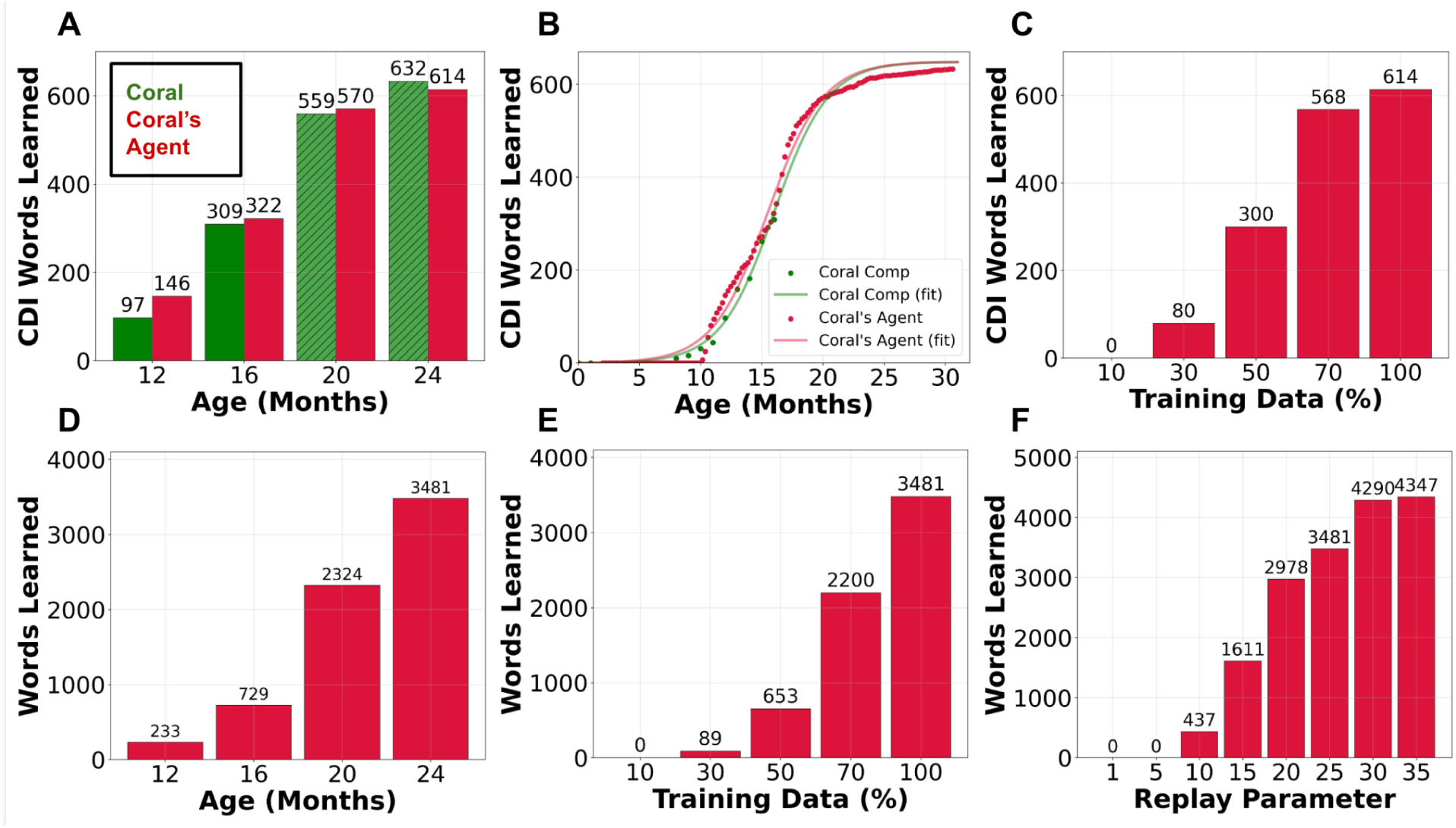
Word-learning trajectories from a single-infant home environment (Coral). (A–C) CDI analysis: (A) Number of CDI words known by Coral from monthly CDI comprehension reports (green solid bars) and extrapolated estimates (green hatched bars), alongside the number of CDI words learned by the model (red) at 12, 16, 20, and 24 months. B) Full CDI word-learning trajectories for Coral (green) and the model (red), including observed data points and sigmoid fits. (C) Effect of daily input quantity: CDI words learned at 24 months as a function of the fraction of daily available data relative to the maximum. (D–F) D) Open-vocabulary analysis: Cumulative learned lemmas over development, E) Effect of training-data fraction at 24 months F) Effect of replay count on word learning. Counts reflect temporally aligned predictions; a word is considered learned once its correct-prediction count exceeds the learning threshold (Methods).

The strong alignment between Coral’s developmental trajectory and that of Coral’s agent would not be possible without the ultra-dense 1kD recordings (Fig. 5-C). To assess how recording density affects learning, we trained the agent on different fractions of the available daily data. Figure 5-C shows the number of CDI words learned by 24 months as a function of recording density. With only 10% of the daily data (slightly less than 1 hour of daily speech), the agent learns no CDI words by 24 months. At 30% (∼2.6 hours/day), it learns only 80 of the 649 words (∼12%). The relationship between data fraction and words learned was nonlinear: at 70% of the daily recordings, Coral’s agent learned roughly 90% of the CDI words achieved with 100% of the data. Still, achieving full alignment between Coral’s and the agent’s learning trajectories required the complete set of recordings, likely because language input has a heavy-tailed distribution. These results demonstrate that ultra-dense naturalistic recordings are essential for training agents on everyday experience in a way that accurately captures real-world language statistics.

Coral’s agent acquired a large vocabulary of thousands of words over 24 months of training. Because the CDI list captures only a subset of the words Coral encountered during the first two years, we estimated the agent’s capacity to learn all words available in the home environment. We counted word types at the lemma level, collapsing inflectional variants (e.g., *eat*, *eating*, *ate*) into a single entry. This analysis pools words across parts of speech; stratifying by part of speech yields the same qualitative pattern. Figure 5-D shows the cumulative number of learned lemmas over development. Because the size of the open vocabulary is not fixed, we report raw cumulative counts rather than fitting a parametric growth curve. As in the CDI analysis, the number of learned words increased steadily with age, reaching thousands of lemmas by 24 months. Vocabulary growth also depended strongly on recording density (Figure 5-E): training on 50% of the daily data (maximum value of 4 hours/day) yielded only about 19% of the lemma vocabulary achieved with the full 8 hours/day by 24 months. This pattern suggests that denser sampling enables the agent to acquire the long tail of low-frequency words not captured in the CDI list.

#### Replay strongly shaped learning outcomes

Augmenting the daily training protocol with resampling of recent events from memory (“replay”) was critical for the agent’s learning (Fig. 5-F). Although Coral’s language environment was very rich - captured by ultra-dense daylong recordings totaling roughly 10 million tokens per year - single-pass exposure to the data was still insufficient for effective learning (Fig. 5-F, 1 exposure). To support learning, we incorporated replay by revisiting past events at the end of each day. Specifically, we resampled prior events in an amount matched to that day’s awake hours, using an inverse weighting function that favored more recent events. The results indicate that repetition is critical: with little or no replay (e.g., 1–5 exposures), the agent struggled to learn words, consistent with the idea that replay-like mechanisms support development and learning^54^. We found that roughly 25–30 total daily exposure cycles were sufficient to support development, with additional replay providing only minimal benefit.

In addition to the number of words learned, we also assessed learning pace as a function of the model’s internal parameter - batch update rate. Figures S-4 plot the sigmoid-fitted time-to-learn parameter (TL) as a function of batch update rate (see Methods). Consistent with the results above, increasing the update rate reduced the time required to learn half of the CDI word list, suggesting that not only the depth but also the frequency of processing incoming experience shapes language acquisition.

#### Generalization across infants

A central goal of developmental psychology is to explain both shared developmental mechanisms and meaningful individual differences. So far, we have shown that the learning agent can simulate a single child’s developmental trajectory in a real-life setting using that child’s ecologically valid input. We next tested whether the agent generalized across children growing up in different home environments. Specifically, we asked whether: 1) we could replicate learning from seven additional sets of recordings from children in distinct home environments; and 2) the framework could account for meaningful individual differences in vocabulary acquisition and learning rates across all eight children.

To examine robustness and generalization, we trained separate learning agents for seven additional children in the 1kD Project, whose everyday experiences were recorded and processed using the same protocol. For each child, a distinct agent was trained on that child’s home recordings, and learning progress was tracked longitudinally using the same analyses described above. As with Coral’s agent, we quantified phoneme and word learning over time and extracted summary parameters by fitting learning curves to each child and the corresponding agent. For five out of the eight children, our ultra-dense recordings were sufficient to support training for approximately 30 months (Figure 1-B). For the remaining three children, however, the available isolated-speech input was more limited, totaling 1,900-3,500 hours, corresponding to approximately 14-20 months of training data. For these infants, we extrapolated the agent’s word-learning trajectory over the missing months using the same sigmoid-based procedure used to estimate children’s comprehension beyond 16 months of age (Fig. 2-A; Methods).

We observe robust generalization of the learning process across all eight children (Figs. 6-7), together with precise sensitivity to individual differences in children’s learning trajectories (Fig. 8). Phoneme learning was highly consistent: across all eight learning agents - each trained on data collected from a single child’s household - an average of 37.5 phonemes (min: 36, max: 38) were acquired within a similar time window (≈12–15 months; Fig. 6). Topaz’s agent was the only one to acquire 36 phonemes due to the smaller available training data.

**Figure 6.**
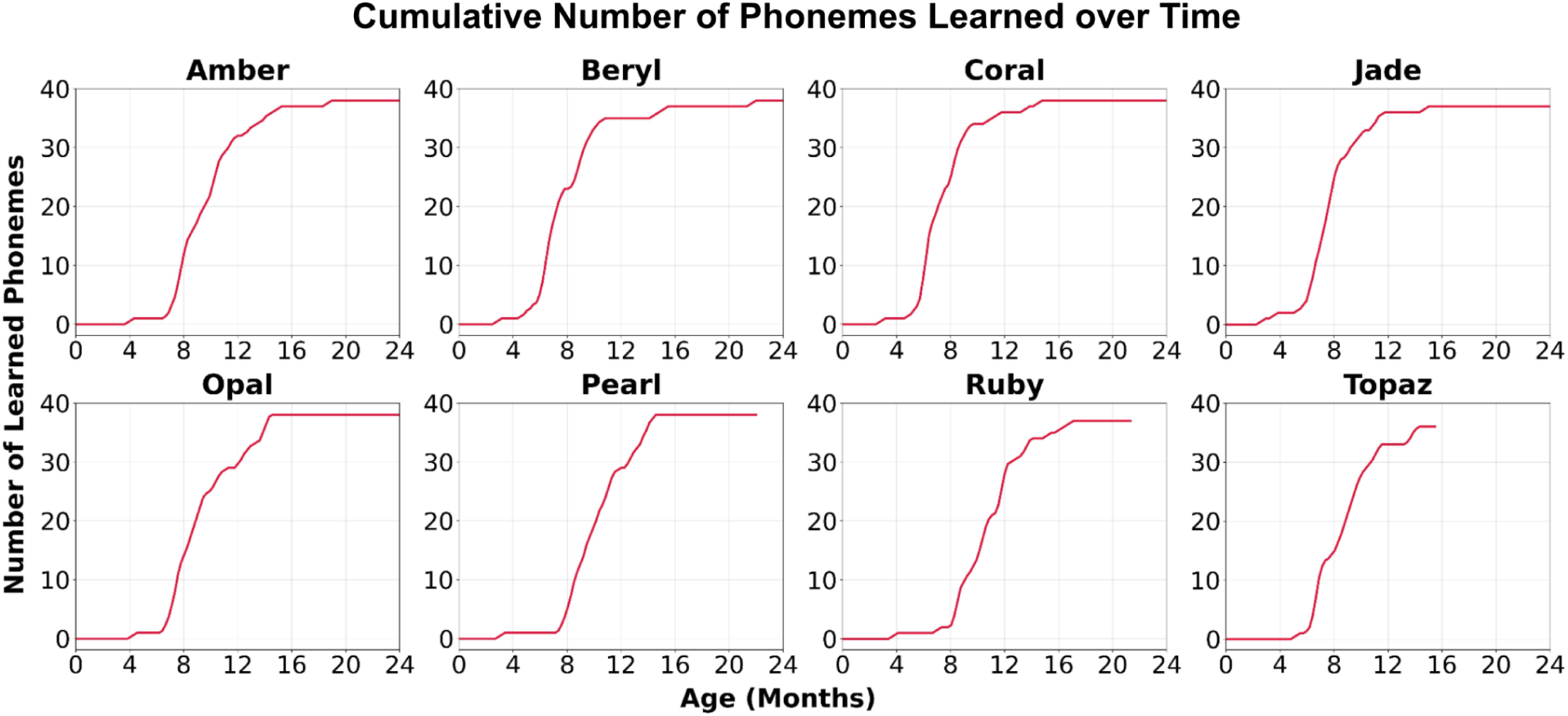
Phoneme acquisition over time across different home environments. Cumulative number of learned phonemes over development using a phoneme-coverage criterion (Methods).

**Figure 7.**
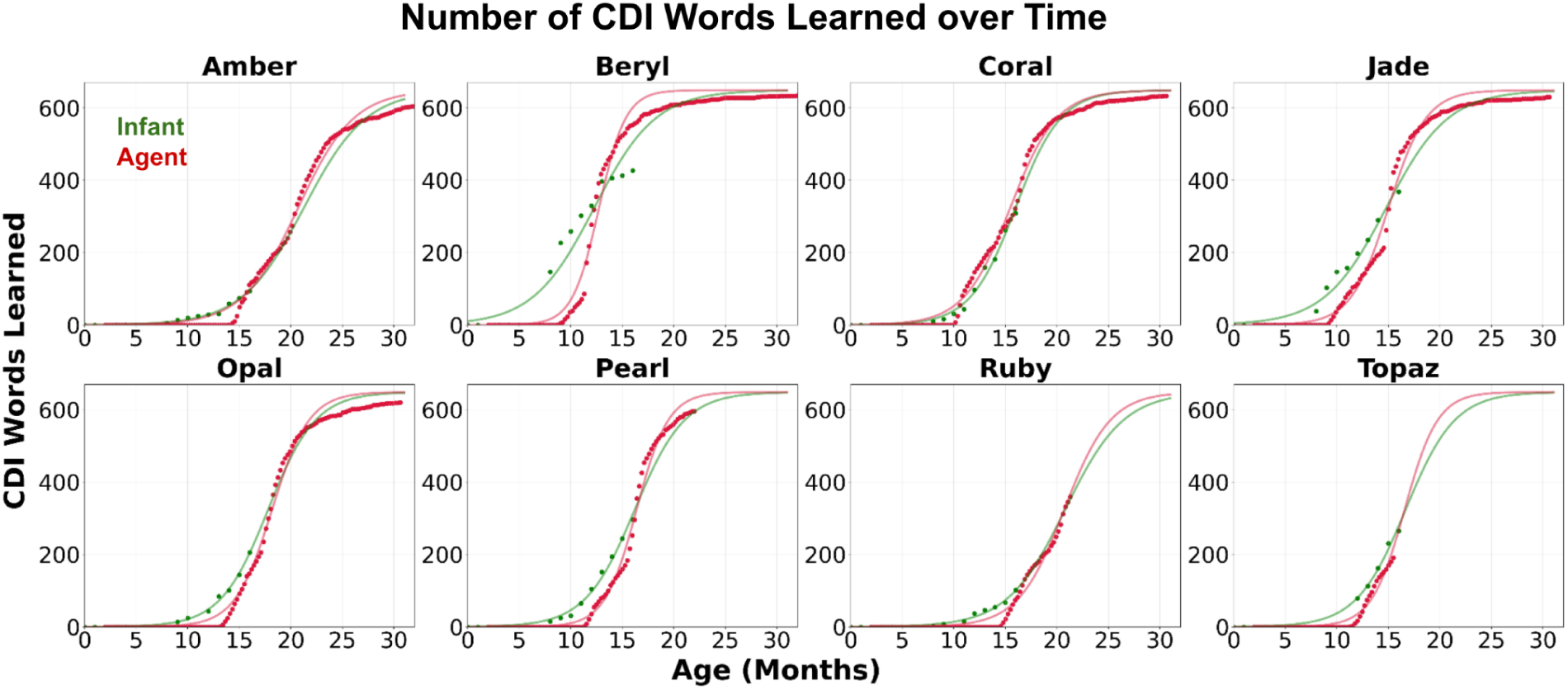
Word-learning trajectories across different home environments. Points show the cumulative number of learned words over development; lines show sigmoid fits to the full trajectories. Green denotes children’s CDI-based comprehension trajectories; red denotes model trajectories.

**Figure 8:**
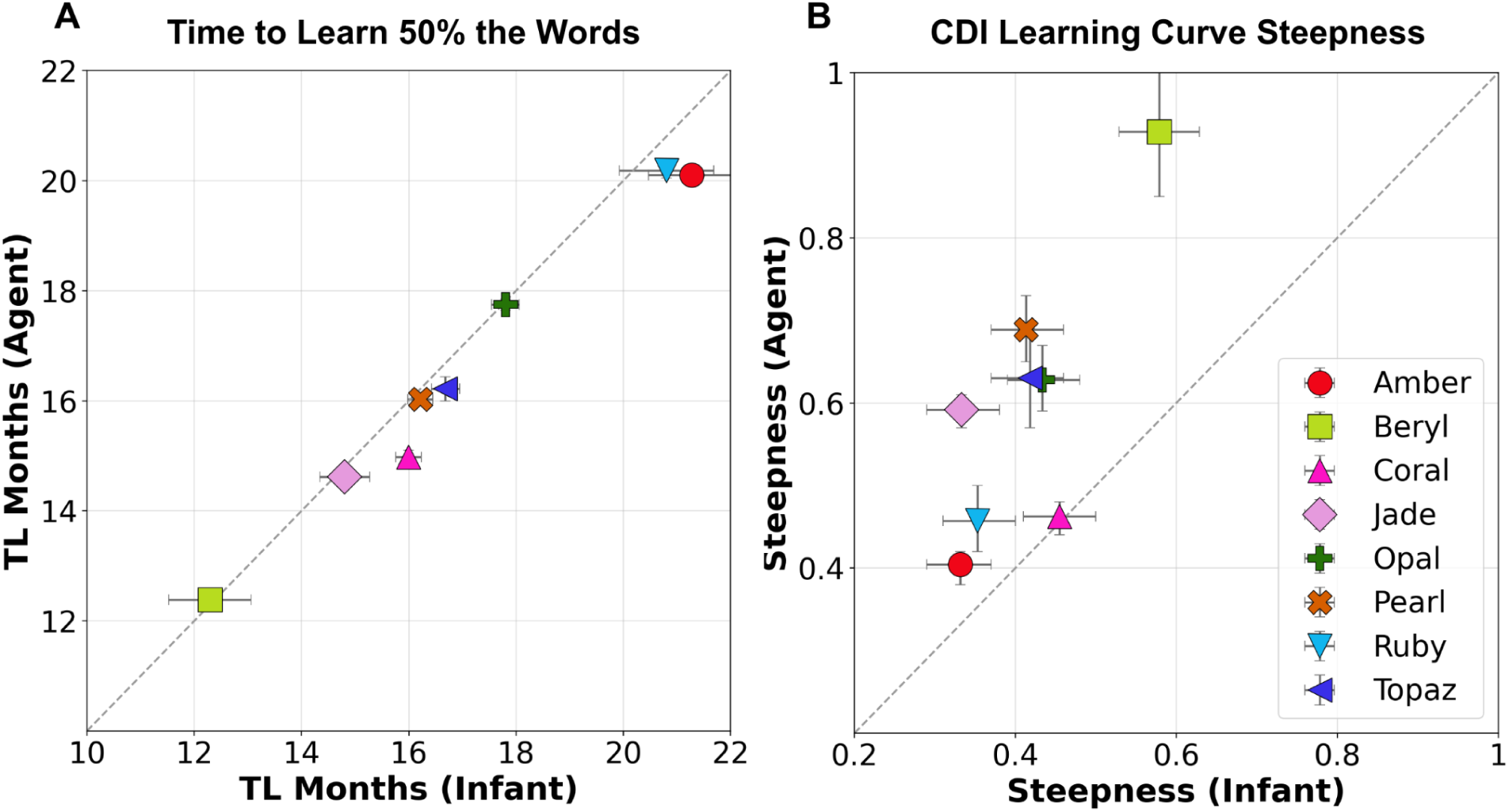
Cross-infant comparison of fitted word-learning trajectories. Scatter plots compare fitted learning-curve parameters for each infant (x-axis) and the corresponding model trained on that infant’s home input (y-axis). (A) Time to reach 50% of words learned. (B) Learning-curve steepness (slope). Each point corresponds to one infant-model pair.

We observed robust word learning across all eight children (Fig. 7). Overall, the agents’ learning trajectories closely matched those of the children, with one notable exception: Beryl exhibited an unusually steep early word-learning curve. Beryl’s CDI report suggests comprehension of more than 150 words by 8 months - an exceptionally high estimate that the sigmoid fit does not accurately capture. In comparison, Beryl’s learning agent reached 150 words at approximately 11 months, and its overall trajectory was otherwise similar to those of the other children. Notably, Beryl’s CDI report also indicated that the child had acquired roughly half of the CDI-list vocabulary (about 325 words) by 12 months, a pattern that is more typical and more closely aligned with the trajectory of Beryl’s learning agent.

To quantify the overall level of agreement across agents and children, we sampled developmental milestones corresponding to 10%–90% of the maximum vocabulary in 10% increments and calculated the mean absolute timing error between each child’s fitted trajectory and their learning agent. Overall, the mean absolute error across the children was 22 days (range 15–56 days).

The learning agent captured individual differences in word-learning rates across all eight children (Fig. 8). To quantify these, we fit a sigmoid function to each child’s learning curve and the corresponding agent’s, and extracted summary parameters. These parameters reflect the time taken to learn half of the CDI words as well as the steepness of the sigmoid curve (the pace of learning). Children differed in the age at which they reached 50% of CDI words, ranging from approximately 12 to 22 months (Fig. 8-A, x-axis). These midpoint ages were closely matched by the estimates from each child’s learning agent (Fig. 8-A, y-axis). Differences in learning steepness were captured only partially (Fig. 80B), largely because the agents exhibited a slightly delayed onset of word learning, followed by a somewhat faster learning rate thereafter.

We also report additional group-level analyses comparing agent and infant learning trajectories. Figure S-5 (supplementary) compares observed and extrapolated vocabulary sizes for infants and their corresponding agents at 12, 16, 20, and 24 months. Across infants, agent estimates closely match infant vocabulary at each age and show the expected monotonic increase over development. Figure S-6 (supplementary) shows the cumulative number of learned dictionary words over time, reaching vocabulary sizes on the order of thousands by ∼30 months for agents with sufficiently long training windows and exhibiting similar growth trajectories across individual agents. Finally, Figure S-7 (supplementary) shows strong agreement in the rank order of nouns acquired during learning, with weaker correlations for pairs with shorter training periods.

## Discussion

Children grow up in complex, dynamic, and highly individualized environments, but nonetheless converge on broadly shared linguistic systems. To explain how such convergence arises from real-world experience, we paired ultra-dense longitudinal recordings of children’s daily auditory input with a learning agent trained on each child’s unique language experience using a cognitively plausible protocol and no prior knowledge of speech or language structure. This provided a mechanistic account of how speech representations and early lexicons can emerge from naturalistic input over developmental timescales. Notably, the agent learned fine-grained speech units that aligned with the English phoneme inventory and subsequently built a lexicon whose growth closely paralleled that of individual children. These results demonstrate that naturalistic environments contain sufficiently rich structure to support early speech perception and language learning when coupled with appropriate learning mechanisms, while also accounting for individual differences in developmental trajectories.

The learning-agent architecture and training protocol provide a mechanistic bridge between early speech processing and lexicon formation. Phoneme-like categories were not presupposed, but instead emerged as regularities across fine-grained 20-ms discrete speech units. Critically, the discrete units were learned directly from environmental speech by an encoder designed to identify reliable structure in the continuous signal. Word-like units, in turn, emerged in the subsequent prediction module (decoder) as consistently predictable sequences of these discrete units.

Importantly, both phoneme- and word-like representations were acquired without explicit language priors or external supervision. This account aligns with mechanistic implementations of self-supervised speech learning^39,55^ and with behavioral evidence of gradual tuning to native-language speech units in the first year of life^13,15^. In the learning agent, as in children, early speech discrimination was associated (by design) with later aspects of language development, such as word comprehension^56^. Finally, the timescales over which phoneme- and word-level representations developed closely matched prior reports^12^ and those observed in our focal infants.

Development unfolds within rich, multidimensional environments that are not accurately captured by sparse measurements. The 1kD recordings provide an unprecedented view of children’s everyday experience, with a density and temporal resolution that far exceed traditional laboratory sampling or brief naturalistic observations. These ultra-dense recordings were essential for training the agent: learning performance degraded substantially when daily speech input fell below 8 hours (Fig. 5C, E). Our findings suggest that early language acquisition depends on two related factors: a rich linguistic environment that supports early speech learning and boosts vocabulary growth, and access to sufficient long-tail statistics that allow the acquisition of infrequently used words, which are typically learned slowly. More broadly, these results imply that modeling naturalistic development requires dense, continuous sampling of language input as it unfolds over minutes and hours and accumulates across weeks, months, and years.

While we found the speech environment to be rich, effective learning also required further enrichment through replay. Without a daily replay of recent input, the agent failed to learn stable speech units and to accumulate lexical structure over developmental timescales. Replay transformed fleeting sensory experiences into consolidated representations that supported long-term learning, paralleling influential theories of memory consolidation in which offline reactivation stabilizes recent experience, reduces interference, and enables gradual abstraction^57–59^. Consistent with this account, replay-like reactivation has been observed during sleep at the neural level in hippocampal ensemble activity, providing biological evidence for offline reinstatement mechanisms that reshape representations over time^60^. In infancy, sleep has likewise been implicated in the reorganization and generalization of newly learned word meanings, supporting the plausibility of replay-like consolidation during early learning^54^. Together, these findings suggest that development depends not only on daily experience but also on iterative cycles of experience and consolidation, with sleep-dependent replay potentially linking perception, memory, and learning during early cognitive development^61,62^.

Our agentic framework provides a novel, data-driven benchmark for developing and evaluating theories of cognitive development. By integrating structured learning architectures with rich, ecologically valid input, researchers can now model how learning unfolds over individual developmental timelines in real-world contexts. Constructs typically studied in brief, tightly controlled laboratory tasks can thus be reformulated as explicit computational hypotheses about how naturalistic input is transformed into developmental outcomes. The framework’s emphasis on mechanistic specificity is particularly powerful: it requires theories to be instantiated as algorithms that learn under the temporal and statistical constraints children face in everyday life, while enabling direct comparisons of their explanatory value against alternative accounts^63–65^.

Several limitations delineate important directions for future work. First, the current agent was trained on clean, resynthesized speech to mitigate background noise and overlapping speakers, which necessarily simplifies the acoustic complexity of everyday input. Second, the present framework omits multimodal information and social context that support children’s learning^66^.

Incorporating biologically plausible mechanisms for speech-nonspeech separation, robust learning under realistic noise and speaker overlap, and audiovisual integration will be essential for ecological completeness. Third, extending the framework beyond early sound and word learning to syntactic and pragmatic learning will test whether similar computational principles scale to capture higher levels of linguistic structures^2^. Finally, developing an embodied reinforcement-learning agent that can act in and learn from its environment could further enrich the modelling framework by linking perception, action, and language learning as it happens in real life.

In conclusion, the 1kD dataset together with the learning agent provides a novel mechanistic account of how rich environmental input (nurture) interacts with internal learning mechanisms (nature) to support early language development in individual children. Furthermore, it demonstrates that foundational aspects of language- phoneme-like and word-like representations- can emerge through adaptive direct fit to the rich structure of young children’s natural environments^67^. More broadly, this agentic modeling approach aligns with views of development is an emergent product of multiscale interactions unfolding over time, rather than the result of fixed, pre-specified representations^65^. The 1kD framework thus offers a concrete foundation for the next generation of empirically grounded, ecologically robust theories of language acquisition.

## Methods

### The First 1,000 Days corpus

This study used data from eight families enrolled in the First 1,000 Days Project (see Raviv et al. for details^25^). Each home contained 4-14 recording devices distributed across 2-5 rooms. Recordings were collected from February 2022 to May 2025, yielding approximately 600,000 hours of audio-video data. Longitudinal behavioral measures were collected alongside the recordings, including caregiver questionnaires and developmental assessments. Our primary measure was the MacArthur-Bates Communicative Development Inventories (CDI^44^), administered monthly from 8 months onward to provide dense within-child measures of language development.

### Corpus construction

Full details of the recording and processing pipeline are provided in Raviv et al.^25^. For each learning agent, we used transcripts of all speech detected in the vicinity of a child across the eight households. Recordings were selected using an AI-based multimodal preprocessing pipeline for child and speech detection, which achieved high overall accuracy (mean F1 = 0.89, precision = 0.82, recall = 0.97).

The raw far-field recordings contained substantial background noise and reverberation. To model learning from linguistic input, we generated a clean speech-only signal while preserving linguistic content and temporal order. All speech produced around each child was first transcribed. The resulting transcripts were splitted into individual sentences using spaCy^68^. Sentences were concatenated into segments to provide sufficient context for synthesis and to better approximate continuous natural speech; segments had a mean length of 34 words.

Each segment was synthesized with a text-to-speech (TTS) model^45^ conditioned on a voice prompt (a short audio sample used to make the synthesized speech sound like a particular speaker). For each household, we used a mixture of eight adult female voices. Thirty percent of segments were synthesized using a voice matched to the child’s mother, and the remaining 70% were distributed evenly across the other voices. Speaker identity for each segment was sampled from this distribution. This procedure preserved the lexical and distributional structure of the input while removing natural speaker-specific variation, prosody, background noise, and reverberation.

#### Quality Control with Transcription Matching

After synthesis, each generated segment was transcribed with WhisperX^69^. The resulting transcript was compared with the original segment text using normalized Levenshtein distance, and the segment was retained only if the distance was below 0.1. If this criterion was not met, synthesis was repeated up to five times. This verification step ensured that the synthesized audio preserved the original linguistic content.

#### Speaker Voice Selection

We isolated the primary caregiver’s voice using one of two approaches. For families with LENA recordings, we extracted speech samples in which the main caregiver was in close proximity to the infant. For the remaining families, we selected high-quality speech samples from our recordings using person-detection metadata together with background-noise estimates to identify clean speech.

#### Final Output

For each segment in the speech corpus, we saved both the generated audio and the associated transcription. This allowed us to later align the speech with the linguistic input and use it for evaluating the developmental learning process.

### CDI list analysis

We counted each word on the CDI checklist in the language corpus using the listed surface form together with common morphological variants (e.g., plural nouns and tense-inflected verbs).

Because the present analyses focused on single-word learning, we excluded two-word phrases from the processed list. We also removed idiosyncratic items, such as the infant’s own name (3 overall), to ensure comparability across children. This yielded a final set of 649 words, corresponding to 95% of the 680 CDI items. We used this maximum vocabulary size (N = 649) when fitting the sigmoid function.

### The Learning Agent Architecture

We used the same learning-agent architecture for all children. Specifically, we implemented a two-module encoder-decoder model to capture the developmental trajectory of children’s word learning (Fig. 3-A ^38,39,70^). The encoder was based on DINO-SR^49^ and closely followed the configuration reported in that work. It consisted of a convolutional feature extractor followed by a Transformer^26,71^ with K=12 layers, 12 attention heads, and a 768-dimensional embedding space.

Audio was sampled at 16 kHz. The student network was trained with sequence masking, in which 80% of input features were masked before the Transformer to encourage robust, context-dependent representations. The codebook contained 512 discrete speech-unit clusters, and encoder outputs were generated at a fixed temporal resolution of 20 ms.

The decoder followed the architecture of Kharitonov et al.^50^. Consecutive identical speech-unit labels extracted from layer 9 of the encoder were collapsed into single tokens and paired with their aggregated durations. The decoder was trained to predict the next speech unit, thereby modeling the input’s sequential structure. It consisted of a Transformer with K=12 layers, 12 attention heads, and a 768-dimensional embedding space.

The encoder and decoder were trained jointly with Adam (β1=0.9, β2=0.98). The encoder used a learning rate of 1×10−4 with a 16,000-step warm-up. The decoder used the same base learning rate, modulated by an iteration-dependent sigmoid schedule parameterized by a transition point, Tp, and transition rate, α=0.0003. Per-family fitting details for Tp, together with the batch sizes used for both modules, are reported in the Training protocol section. Gradients were backpropagated separately through the encoder and decoder. The total training objective combined two terms: masked cluster prediction and next-unit prediction, each weighted 1 and 10, respectively.

Each agent model was trained on a single NVIDIA H100 GPU. In a typical setup (X = 25 see below, total number of training months = 30), training took 42 hours, while the time-dependent analysis took approximately 10 hours.

### Training Protocol

#### Daily speech input parameter

Our environmental language corpus included detailed logs of the minutes during which the infant was detected in each household, along with the number of words spoken in each minute (see Raviv et al., for details^25^). To estimate the number of words spoken per hour for each infant, we identified all 20-min intervals of continuous speech across recording days in each home. For each interval, we computed the mean number of words per minute and converted it to an hourly rate by multiplying by 60. We then averaged these hourly rates across days and hours to obtain a single household-level estimate of words per hour. The resulting values ranged from 2670 to 3806 words per hour (median = 3520.5).

To account for the developmental increase in awake time, we defined a monotonically increasing function, *F*(*d*), that specifies the number of awake hours with available speech per day at each age. This function was intended to approximate the age-related increase in infants’ effective awake and attentive time and was based on estimates of the maximum number of words heard per day^30,31^. We assumed that *F*(*d*) was shared across families and increased from 2 hours per day in the early months to 8 hours per day by 14 months of age, remaining constant thereafter (Fig. 3-C).

For simplicity, we assigned the same value *F*(*d*) to all days within a given month and assumed 30 days per month in subsequent calculations. Because early speech detection and its separation from background sounds were not modeled here^38,39^, training began at 2 months of age.

#### Batch size parameter

To align the agent’s learning time with each infant’s developmental timeline, we fit the batch size (BS) parameter separately for each infant using CDI-derived learning milestones. For each infant, we fit a sigmoid function to the CDI comprehension trajectory (Results, Section 1; Figure 2-A) and extracted *D*_50_ the age in days at which the model predicted comprehension of 50 words^1^. We then converted this milestone into cumulative speech exposure by integrating the age-dependent speech-hours schedule from 60 days (2 months) up to *D*_50_ assuming 30 days per month. The lower bound of 60 days reflected that the agent model does not include early speech detection. Cumulative exposure at the milestone was defined as:

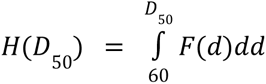 *F*(*d*)*dd*, where *F*(*d*) is measured in hours/day, so the integral gives total hours of speech exposure.

To relate developmental time to training time, we defined an updates-per-hour rate, *U*_*h*_, which links real-world exposure to optimization steps. For a setting with X daily passes over the data (see replay details below), the total number of optimization steps accumulated by the milestone *D*_50_ is:

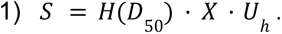

We selected an infant-specific value of *U*_*h*_ such that the implied number of steps at the 50-word comprehension milestone was on the order of the number required for encoder stabilization (∼10^5^ steps). Thus, *U*_*h*_ ∼ 100000/(*H*(*D*_50_) · *X*). This procedure yielded one update-per-hour value for each infant. Finally, we derived the batch size by dividing by the total number of files per hour (*W*_*h*_/34, 34 is the average number of words per segment) by the number of updates per hour (34)

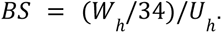

#### Replay parameter

Our replay mechanism simulated daily re-exposure by resampling previously encountered data after a single pass through the current day’s speech. The sampled events were then replayed X−1 additional times, where X denotes the total number of exposures, including the current day’s input. The amount of replayed speech was set to match the number of awake hours containing speech on that day. Replay events were generated using an inverse sampling distribution, where the likelihood of selecting a specific day decreased as a function of temporal distance. The repetition parameter, X, was selected from {25, 30, 35, 40} and fit separately for each infant to best align the agent’s learning trajectory with the infant’s observed developmental trajectory.

#### Decoder learning rate function

The sigmoid transition point, Tp (Fig. 3 B-3), was fit separately for each infant to align agent time with developmental time. We set Tp such that the agent’s effective learning time at the transition corresponded to the age at which the infant was estimated to have acquired approximately half of the CDI target words. Following the procedure described above, we (i) obtained this milestone age from the infant-specific CDI sigmoid fit (*D*_*h*_), (ii) converted it to cumulative speech exposure in hours using the age-dependent speech-hours schedule, *H*(*D*_*h*_) and (iii) mapped this exposure to optimization steps, S, using Eq. 1 together with the previously estimated updates-per-hour rate.

### Evaluation

To match real-world learning conditions, the learning agent was trained directly on audio rather than text. To evaluate learning at the phoneme and word levels, we used the text transcripts associated with each audio file. Each audio file was aligned to its transcript with the Montreal Forced Aligner (MFA), run with default settings and the english_us_arpa pronunciation set^72^. After collapsing stress variants, this procedure yielded 39 phoneme categories. MFA provided time-stamped annotations for the onset and offset of each word and phoneme.

### Phoneme Learning

#### Mapping phoneme tokens to discrete speech units (dominant unit per token)

The encoder outputs a time series of discrete speech-unit identifiers, which we collapse into unit labels with associated durations. For each phoneme instance, *p i* we considered all units whose timestamps fell within the phoneme interval and defined the dominant speech unit, *u*_*i*_, as the unit with the greatest total duration within that interval. We used the dominant unit to avoid overweighting longer phonemes and to reduce the influence of boundary frames, where coarticulation and acoustic transitions increase label variability. This procedure yielded one pair, (*p*_*i*_, *u*_*i*_), for each phoneme instance and provided a stable, interpretable mapping between learned units and phoneme categories.

Using these pairs, we constructed a contingency table of token counts, *C*(*p*, *u*) where each entry denotes the number of phoneme instances of type p whose dominant speech unit was u. We quantified phoneme purity, 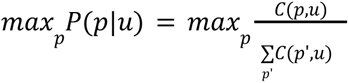 which measures how selectively a speech unit corresponds to a single phoneme. For visualization, we used the phoneme-to-speech-unit association matrix with entries *P*(*p*|*u*) and reordered units according to the phoneme for which each unit attained its maximum value (Fig. 4-A).

#### Learned phonemes via a coverage criterion

To examine phoneme acquisition over time, we computed, at each evaluation time point t, the phoneme-unit count matrix *C*_*t*_(*p*, *u*) and the corresponding phoneme-purity matrix *P*_*t*_(*p*|*u*). Given a purity threshold, τ, we defined the set of high-purity speech units for phoneme p at time t as: *U*_*t*_(*p*; τ) = {*u*: *p*_*t*_(*p*|*u*) ≥ τ}.

The coverage of phoneme p at time t was defined as the fraction of phoneme instances accounted for by these high-purity units. A phoneme was considered learned at the earliest time point t for which (i) coverage exceeded a threshold, γ and (ii) the total number of observed instances of p exceeded a minimum threshold, N. We then plotted the cumulative number of learned phonemes over time (Fig. 4-B). We used τ = 0. 5, γ = 0. 3, *N* = 50 .

### Word Learning

#### Audio-text-model output alignment

The encoder consisted of teacher and student modules (see Results, Section 2). The teacher discretized the audio into speech-unit IDs at a temporal resolution of 20 ms per step (codebook size, 512), and the student predicted these unit IDs with associated probabilities. To form the decoder input, we merged consecutive 20-ms steps assigned the same unit ID into a single token and recorded its duration as the number of merged steps. The decoder language model then predicted a probability for each token given the preceding context. For evaluation, each audio file was represented as a sequence of these collapsed unit tokens, annotated with start time, duration, mean student encoder probability (averaged across merged steps), and decoder-predicted probability.

#### Alignment of model output with text

We aligned collapsed unit tokens to the transcript using an overlap-based criterion: each unit token was paired with any phoneme or word whose annotated time span intersected the token’s time span. For each aligned pair, we computed the duration of the temporal intersection and normalized it by the duration of the phoneme or word to obtain a fractional alignment score (phoneme or word coverage). This score quantified the extent to which a given phoneme or word was covered by each unit token.

#### Decoder prediction probabilities smoothing

To reduce noise in the decoder prediction probabilities, we smoothed them over time within each audio file using a mean-weighted running average. Each unit token was assigned a weight proportional to its mean encoder probability multiplied by its duration, so longer segments with higher encoder certainty had more influence than short or uncertain tokens. The output probability at each step was a weighted sum of the current probability and the past smoothed probability, with the latter down-weighted by a fixed decay factor.

Formally, let *p*_*t*_ be the decoder probability for unit token t, *w*_*t*_ its weight, α the decay factor, 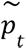 the smoothed probability for unit token t, and *W*_*t*_ and the effective past weight. We initialized 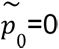 =0 and *W*_*t*_ = 0 calculated iteratively:

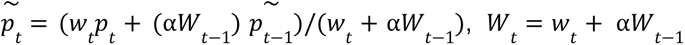

This produced a smoothed prediction probability for each unit token, which was used in the following steps.

#### Word segmentation and learning evaluation

We used the decoder’s smoothed prediction probabilities to track the emergence of word knowledge. Our working assumption was that a word form had been learned when the model could predict a continuous sequence of unit tokens with consistently high confidence in context. To operationalize this, we segmented the unit-token stream using the following rule: a new predicted segment was initiated whenever (i) a transcript-derived word onset was reached, (ii) the smoothed probability of the current unit token fell below an error threshold (mistake score, 0.05), or (iii) the smoothed probability dropped sharply relative to the maximum value observed earlier in the current segment (drop >0.3) while the absolute probability was also low (<0.6). Extracted segments were then matched to transcript word intervals using temporal overlap. A word was counted as correctly predicted at a given time point if at least 90% of its annotated duration overlapped with a predicted segment. To ensure that the evaluation was context-dependent, we excluded the first second of each recording segment (skipped the associated units).

We applied this procedure to each held-out evaluation split, with each split corresponding to one week of data in chronological order. For each week, we updated a growing vocabulary (the child’s “dictionary”; Fig. 3-A) by accumulating correctly predicted words. Each dictionary entry stored the word’s surface form, part-of-speech tag, lemma (assigned with spaCy from the transcript), and constituent phonemes. For learning analyses, lexical types were defined as groups of inflectional variants sharing the same lemma (e.g., *eat*, *eating*, and *ate*). For each lemma, we tracked the cumulative number of correct predictions across weeks. The inflectional variants used for counting CDI words were taken from a predefined list of common morphological variants (e.g., plural nouns and tense-inflected verbs) for each word (see CDI list analysis above). A word was considered learned once its cumulative correct-prediction count exceeded a fixed threshold (N=15).

## Supporting information

Full Supplementry

## Acknowledgements and funding

We are deeply grateful to the 1kD families for their extraordinary generosity and participation. We thank Irene Kopaliani, John Wiggins, and Mark Ratliff for the exceptional institutional support. We also thank Marian Faryna, Stanislav Samko, and Olga Chikhiro of Sigma Software, and Ricky Beaty and Dean Johnson of Beaty Consultancy, for their contributions to the project’s software infrastructure. We are especially grateful to Chandra Greenberg for her work with participating families. This work was supported by Wellcome Leap, the McGregor Girand Charitable Endowment, Princeton Language and Intelligence, Princeton Precision Health, Data-Driven Social Science, and the AI Lab. The content of this publication and any related materials is solely the responsibility of the authors and does not necessarily represent the official views of the funding agencies.

## Authors contributions

H.R and U.H devised the modeling framework and analysis plan, L.H, C.L.W devised the ethical and behavioral data collection framework, H.R, U.H wrote the paper, L.H and C.L.W critically revised the paper, L.H, C.L.W, K.G and A.A performed human data collection and analysis; H.R., L.H, B.C., H.W. performed data collection. A.T, H.R, performed modelling framework implementation, models training and analysis. O.R. and T.R. served as consultants on the modeling framework, advising on model selection, infrastructure, coding, and data collection decisions.

## Footnotes

1 For Baryl whose sigmoid fit yielded implausible estimates, we used the minimal measured value - 8 months.

