## Supplementary material for "The First 1000 Days: An Agent-Based Model of Early Language Acquisition": Full Supplementry

### Supplementary Materials

### 1 - Phoneme Learning Analysis

**Phoneme - speech-unit associations:** We quantified the relation between discrete encoder speech units and phonetic categories over developmental time using purity metrics derived from the phoneme-unit co-occurrence table, following prior work (for example, DINO-SR[46]). As described in the Methods, each phoneme instance p was aligned to the audio timeline and paired with its dominant encoder unit u, defined as the unit occupying the largest fraction of the phoneme's duration, yielding one (p,u) pair per phoneme instance. From the resulting count table, C(p,u), we computed two directional purity measures. Phoneme purity quantified how selectively each unit mapped to a dominant phoneme, $max_p P(p|u) = max_p \frac{C(p,u)}{\sum_{p'} C(p',u)}$, whereas cluster purity quantified how consistently each phoneme mapped to a dominant unit, $max_u P(u|p) = max_u \frac{C(p,u)}{\sum_{u'} C(p,u')}$. For summary curves, we report token-weighted averages of these quantities using unit frequencies for phoneme purity and phoneme frequencies for cluster purity. Across development, these measures showed that the learned speech-unit inventory became increasingly informative about phoneme identity without converging to a one-to-one phoneme code. As shown in Fig. S1-A, phoneme purity increased over training and stabilized at a level consistent with high specialization of speech units to phonetic categories (≈0.6). By contrast, cluster purity remained low throughout training (Fig. S1-B; ~0.14 at stabilization), indicating that individual phonemes were distributed across multiple units rather than mapped to a single dominant unit. This pattern is qualitatively consistent with prior evaluations of DINO-SR[46].

**Within-token dynamics of speech-unit assignments:** To characterize the dynamics underlying the phoneme - unit mapping, we quantified within-token variability using the full sequence of speech-unit assignments within each phoneme segment. For each phoneme token, we computed (i) the number of distinct units observed within the segment and (ii) the fraction of frames assigned to the most frequent unit (dominant-unit share). Figure S1-C shows that within-phoneme unit diversity increased rapidly early in training, peaking at approximately 3.8 units per token, and then declined steadily to approximately 2.5 as learning stabilized. The dominant-unit share showed the complementary pattern (Fig. S1-D). Together, these results indicate that the learned units formed a rich, context-sensitive partition of acoustic space that became progressively more informative about phoneme identity while retaining substantial sub-phonemic structure, consistent with the view that phonetic learning does not require the direct acquisition of discrete phoneme categories[33].

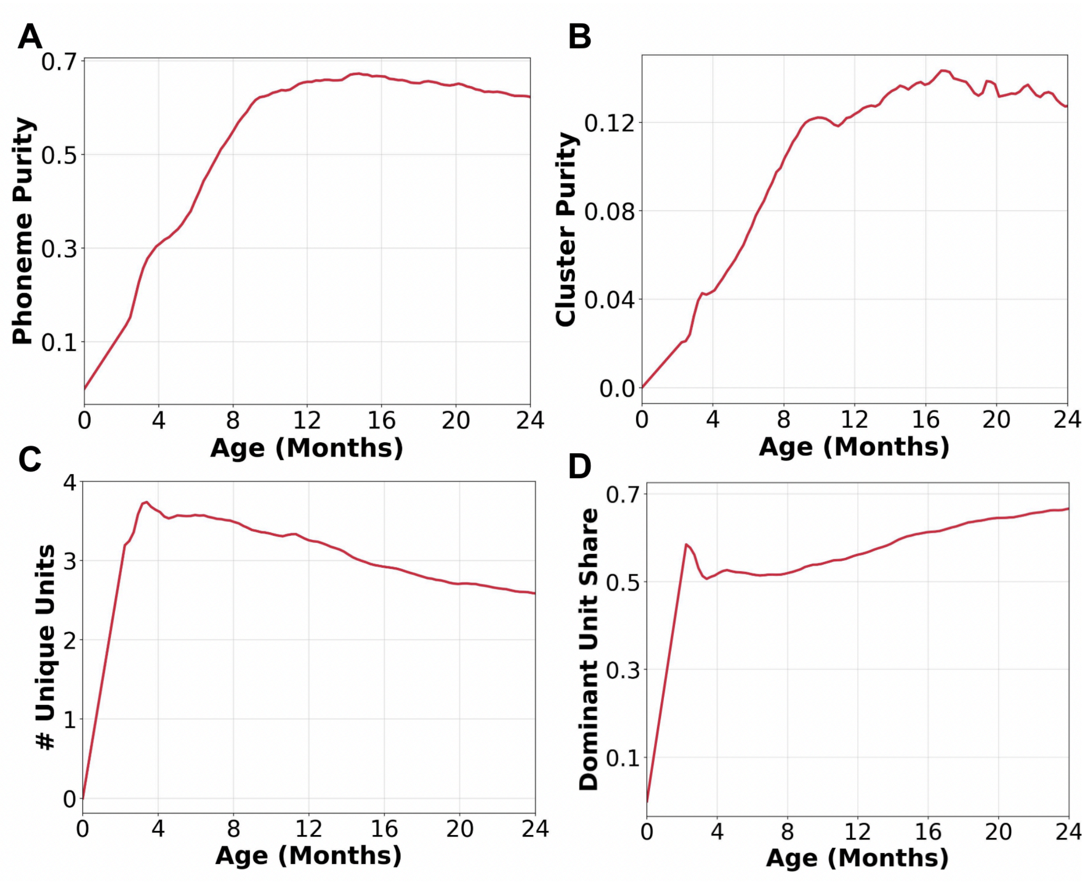


**Figure S1: Phoneme - speech unit alignment and within-token consolidation over time (Coral's agent).** (A) Phoneme purity (unit->phoneme): how selectively each unit corresponds to a single phoneme, summarized over units (token-weighted). (B) Cluster purity (phoneme->unit): how consistently each phoneme maps to a single dominant unit, summarized over phonemes (token-weighted). (C) Within-phoneme token unit variation: the average number of distinct units observed within a forced-aligned phoneme. (D) Within-phoneme token unit stability: the average fraction of segment duration assigned to the most frequent speech unit within a phoneme segment.

### 2 - Effect of Word Learning Analysis Parameters

Figures S2 and S3 each contain 12 panels summarizing the trajectory sensitivity analysis (of Coral's learning agent) across parameter settings. Panels are organized by the three analysis parameters listed in the panel titles, ordered left-to-right as **coverage**, **mistake score**, and **skipped units** (see Methods, segmentation rule). Figures S2 and S3 differ only in the acquisition threshold size (N=15 vs. N=20, respectively), i.e., the number of correctly predicted word occurrences required to count a word as learned. Across all parameter and threshold combinations, the resulting CDI learning trajectories were qualitatively similar: in all cases the agent learned more than ~600 CDI words, and the overall trajectory shapes and slopes were comparable, indicating that our conclusions are robust for a range of parameter values. As expected, increasing N yields fewer learned words (a stricter criterion), whereas lower word coverage (e.g., 0.8) tended to yield earlier estimated learning, consistent with a more permissive matching threshold. We adopted a coverage threshold of 0.9 to allow lemma-level variability while remaining conservative, requiring near-complete temporal overlap with the target word interval.

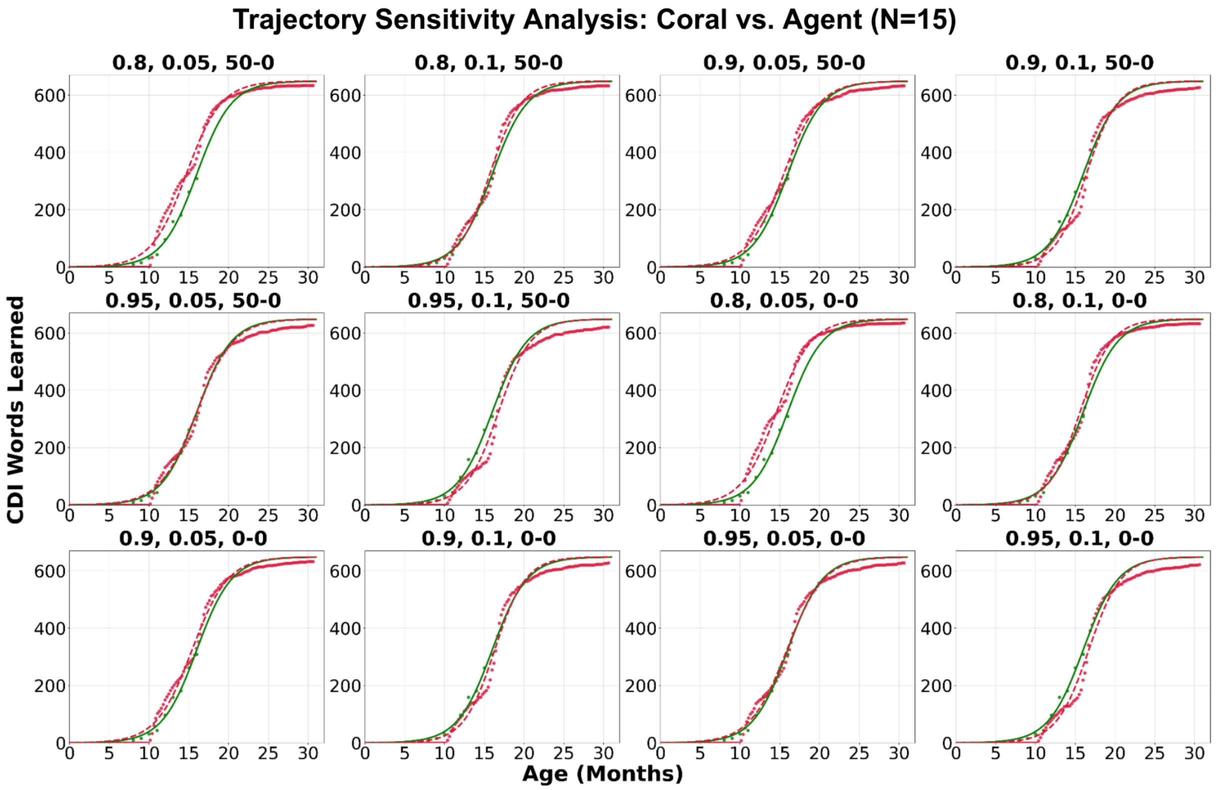


**Figure S2. Trajectory sensitivity analysis (threshold N=15).** CDI learning trajectories for Coral's agent using 12 evaluation settings. Panels are organized by the analysis parameters shown in the panel titles (left-to-right: coverage, mistake score, skipped units; see Methods). A word is counted as learned once it reaches N=15 correctly predicted occurrences.

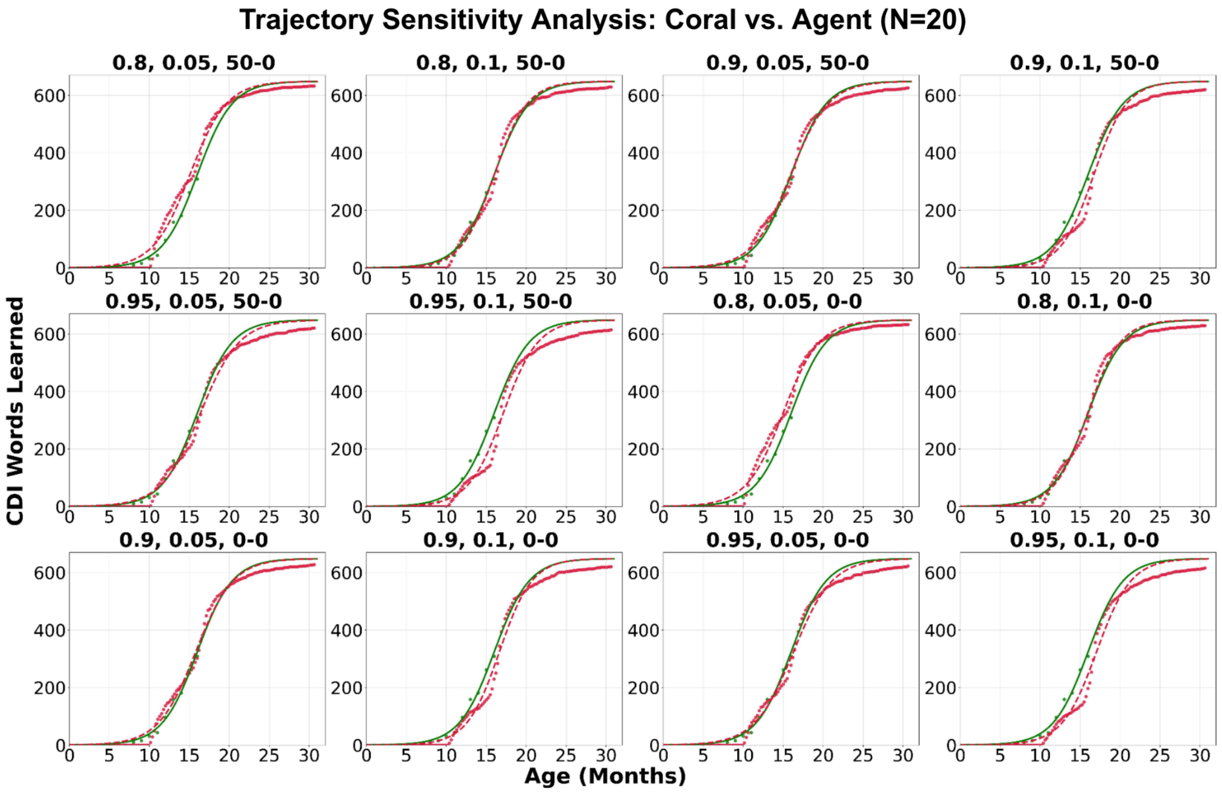


**Figure S3. Trajectory sensitivity analysis (threshold N=20).** Same analysis as Fig. S2, but using a stricter learning criterion (N=20).

### 3 - Changing the hourly model update parameter

Figure S4 shows the learning time of Coral's agent as a function of the updates-per-hour rate, $U_h$, with all other model parameters held constant. For each value of $U_h$, we generated a CDI learning trajectory, fit a sigmoid function as in the main text (Fig. 2), and extracted TL, defined as the age at which the fitted curve reached 50% of its asymptote. We then plotted TL as a function of $U_h$.

Increasing $U_h$ systematically accelerated learning, reducing the time required to reach the milestones captured by the fitted learning trajectory. Thus, $U_h$ can be interpreted as the rate at which the agent incorporates new information per hour of exposure.

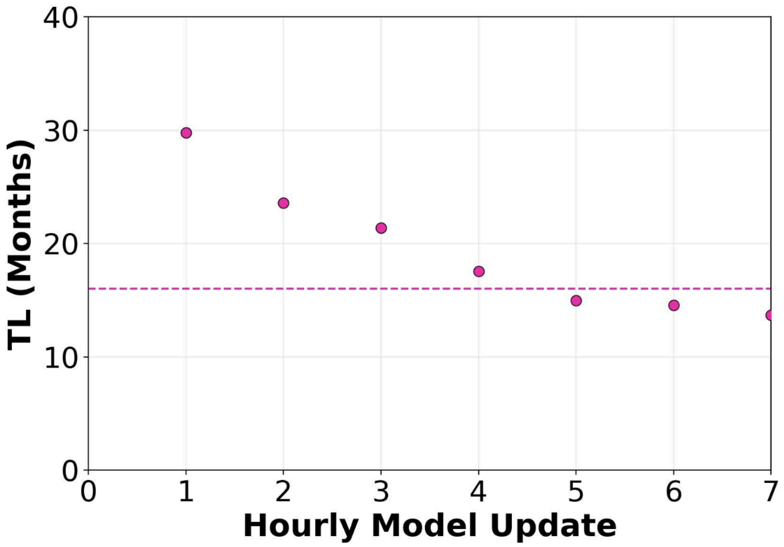


**Figure S4: Effect of agent parameters on learning speed (Coral).** Time to learn half of the CDI word set (TL) as a function of the hourly model update parameter.

### 4 - Group-level results across infants and agents

**Extrapolated CDI vocabulary size:** Figure S5 compares measured and extrapolated CDI vocabulary sizes for infants and their corresponding agents at 12, 16, 20, and 24 months. Using the same procedure as in the main text (Fig. 5-A), we plot observed values when available (solid green bars for infants and solid red bars for agents) and use sigmoid-fit estimates for months with missing observations (hatched light bars). Across infants, agent estimates closely match infant vocabulary at each age and exhibit the expected monotonic increase over development, indicating that the fitted agent trajectories capture both the overall scale and growth of CDI vocabulary across the cohort.

**Dictionary growth trajectories:** Figure S6 shows the cumulative number of learned dictionary words over time. Across agents, the trajectories exhibited a common two-phase pattern, with an initial period of relatively slow growth followed by a steeper increase. For agents with longer training runs, vocabulary sizes reached an order of thousands of dictionary words by approximately 30 months, suggesting continued accumulation of word knowledge over extended training.

**Rank-order agreement in word acquisition:** Figure S7 evaluates whether agents learned words in a similar order to infants by correlating the rank order of word acquisition between each infant and its agent using the noun subset of the CDI. We used infants' production CDI data for this analysis because comprehension data contained too few time points. For agents with sufficient training duration to support age-matched comparisons, correlations were consistently positive and typically exceeded r≈0.40, indicating substantial agreement in acquisition order across the cohort. Three children (Pearl, Ruby and Topaz) had substantially less training data, which limited the age range available for comparison and yielded lower correlations.

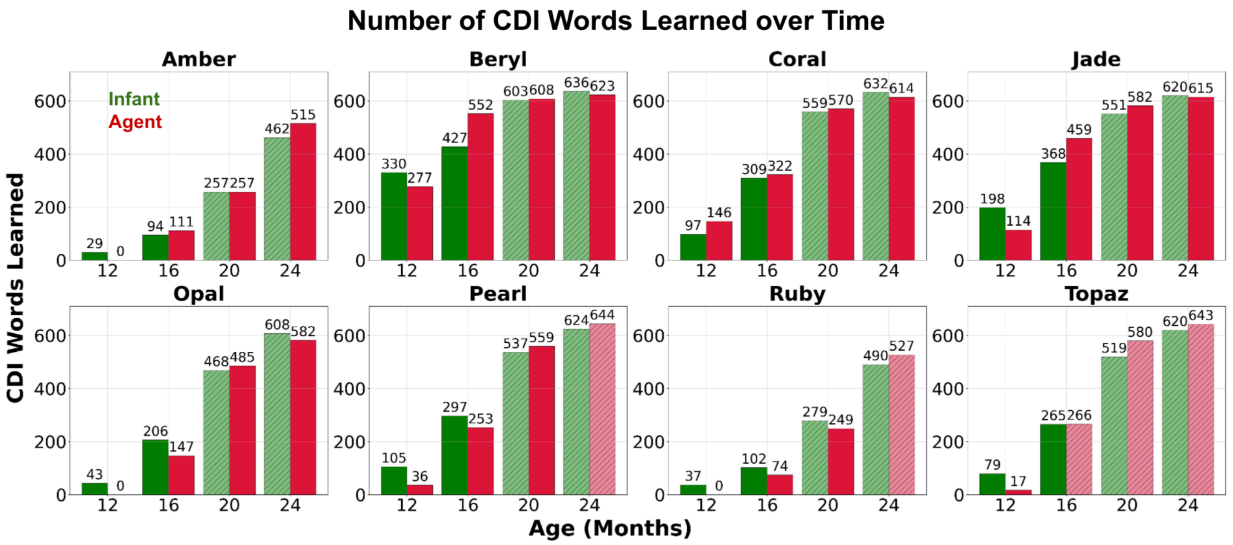


**Figure S5. Extrapolated CDI vocabulary size across infants and agents.** CDI vocabulary at 12, 16, 20, and 24 months for infants (green) and corresponding agents (red). Values are taken from observed measurements when available (solid bars) and otherwise from sigmoid-fit estimates of CDI comprehension trajectories (hatched light bars; Methods).

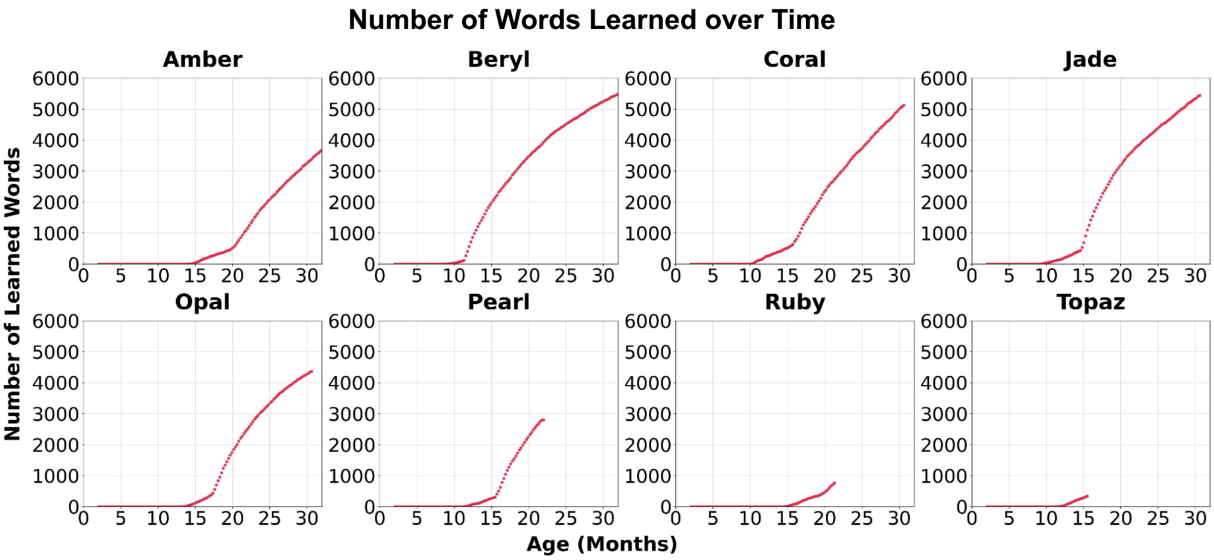


**Figure S6. Dictionary vocabulary growth trajectories across agents.** Cumulative number of learned dictionary words as a function of age/training time for each agent.

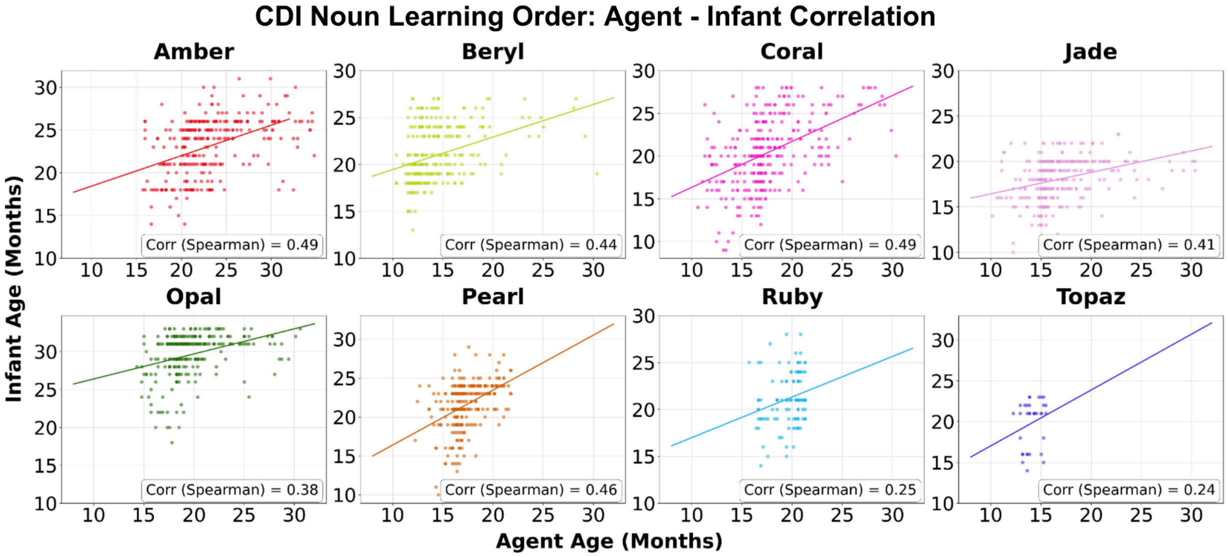


**Figure S7. Rank-order correlation of word acquisition between infants and agents.** Correlation between infants and agents in the order of word learning (computed over Nouns and aligned developmental ages; Methods).
